# Stimulus-specific prediction error neurons in mouse auditory cortex

**DOI:** 10.1101/2023.01.06.523032

**Authors:** Nicholas J. Audette, David M. Schneider

## Abstract

Comparing expectation with experience is an important neural computation performed throughout the brain and is a hallmark of predictive processing. Experiments that alter the sensory outcome of an animal’s behavior reveal enhanced neural responses to unexpected self-generated stimuli, indicating that populations of neurons in sensory cortex may reflect prediction errors – mismatches between expectation and experience. However, enhanced neural responses to self-generated stimuli could also arise through non-predictive mechanisms, such as the movement-based facilitation of a neuron’s inherent sound responses. If sensory prediction error neurons exist in sensory cortex, it is unknown whether they manifest as general error responses, or respond with specificity to errors in distinct stimulus dimensions. To answer these questions, we trained mice to expect the outcome of a simple sound-generating behavior and recorded auditory cortex activity as mice heard either the expected sound or sounds that deviated from expectation in one of multiple distinct dimensions. Our data reveal that the auditory cortex learns to suppress responses to self-generated sounds along multiple acoustic dimensions simultaneously. We identify a distinct population of auditory cortex neurons that are not responsive to passive sounds or to the expected sound but that explicitly encode prediction errors. These prediction error neurons are abundant only in animals with a learned motor-sensory expectation, and encode one or two specific violations rather than a generic error signal.

## Introduction

Sensory responses in the cerebral cortex are influenced by an animal’s behavior^1–10^ and can reflect an expectation for the sensory consequences of movement^11,12,21,22,13–20^. This dynamism is consistent with the theory of predictive processing, which posits that cortical activity prioritizes representing deviations from expectation over directly representing features of the external world^23,24^. Indeed, experiments that alter the sensory outcome of an animal’s behavior have consistently found that neural responses to unexpected outcomes are enhanced relative to expected outcomes^11,16,17,20,22^. This finding may be attributed in part to the recruitment of neurons that are active only when an animal experiences unexpected self-generated outcomes (e.g. ‘prediction error’ neurons)^11,20,22^. Prediction error neurons are believed to represent the violation of a learned sensory-motor expectation, but could alternatively arise from the mixing of movement and sensation signals in a way that is unrelated to expectation^25^. If prediction error neurons are driven by expectation violation, it remains unknown whether they encode a generic error signal or whether they reflect the identity of the unexpected stimulus. Most previous experiments have intentionally violated expectations using a small number of stimuli, and typically only in animals that experienced sensorimotor coupling, making it difficult to distinguish among these possibilities. Here, we used a simple sound-generating behavior in mice to establish the presence of stimulus-specific prediction error neurons with features consistent with a cortical origin for predictive computations.

## Results

### Motor-sensory predictions are specific across multiple acoustic dimensions

The auditory cortex predicts the frequency of a self-generated sound and its expected position within an ongoing movement (Audette et al., 2022, others). But sounds have many features, including spatial location, intensity, and spectral purity. We therefore aimed to determine whether movement-based predictions in the auditory cortex show specificity along multiple acoustic dimensions simultaneously. We trained head-fixed mice to produce a simple sound-generating behavior during which we could precisely control the acoustic outcome of each movement^11^. Mice pushed a lever past a fixed threshold to trigger a water reward (on 25% of trials) when the lever was returned to the home position (Fig 1A). During training, a pure tone (8 kHz) was presented at a consistent position early in each movement, and mice were free to initiate trials ad libitum. Mice rapidly learned to perform the task and averaged more than 2000 sound-generating trials per session.

**Figure 1:**
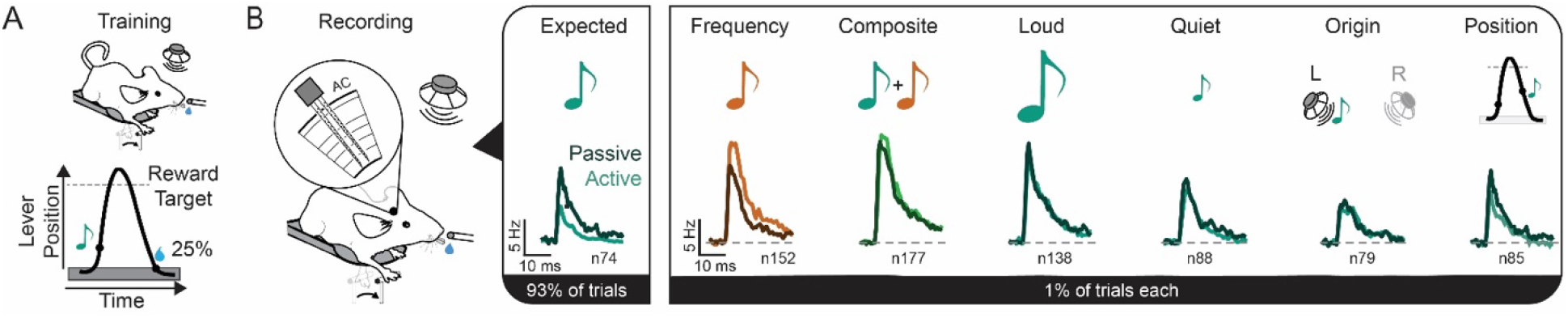
Specific suppression of expected sounds across multiple acoustic dimensions. (A) Schematic of head-fixed lever press training paradigm (Top) and stimulus and reward timing for lever movements (Bottom). Grey area indicates home position. (B) Schematic of multi-array recording sessions in trained mice (Left) and aggregate neural responses to expected and multiple unexpected sounds in the passive (Darker) and movement-evoked (Lighter) context. Of the 1016 regular-spiking neurons we recorded (N = 5 Animals), a subset of neurons are analyzed for each sound type if they respond to that sound in either context (p < 0.01, 0 -60ms post sound onset). Values are listed below each PSTH. Color differences represent sound frequency, and the likelihood of each lever press producing a given sound type during the recording session is displayed in black bar.

Following 10-12 days of training with the lever producing a predictable self-generated sound, we made large channel-count electrophysiological recordings from the auditory cortex while mice executed the learned lever behavior and heard either the expected sound (93% of trials) or a sound that unexpectedly varied in one of several different acoustic dimensions (probe trials, 1% each) (Fig 1B). On these probe trials we did one of the following: substituted a sound shifted 1.1 octaves from the expected sound (Frequency), played an unexpected frequency simultaneously with the expected sound (Composite), changed the intensity of the expected sound by +/-15 dB (Quiet or Loud), changed the spatial origin of the sound (Origin), played the expected sound at the wrong lever position (Position), or omitted the sound altogether. Each of these sounds was also played in a passive listening context during which the lever was removed from the animal’s reach. In total, we recorded from 1016 regular spiking neurons across five animals.

In the passive listening condition, we observed strong neural responses to each sound, including the expected sound (Fig 1B). In the self-generated condition, neural responses to the expected sound were strongly suppressed (∼50%) compared to the same sound heard passively^11^. This strong suppression of neural responses to an expected self-generated sound provides a benchmark for comparing neural responses to unexpected self-generated sounds. If neural responses to an unexpected sound are less suppressed, unsuppressed, or enhanced, we can conclude that the auditory cortex recognizes that sound as a violation of its expectation.

We found that the auditory cortex did not display strong suppression of neural responses to any unexpected sound that we tested. Population-averaged neural responses to the unexpected probe sounds were not suppressed at all (Quiet, Loud, Origin), were mildly suppressed (Position), or were enhanced relative to the passive listening condition (Frequency). As a striking example, we found that neural responses to an unexpectedly quiet self-generated tone were significantly stronger than were responses to the self-generated tone heard at the expected volume. This is in direct contrast to the passive listening condition, during which the louder tone evoked stronger responses than the quieter tone, as would be expected from typical auditory cortex neurons^26^. Importantly, most of the probe sounds used here were of the exact same frequency as the expected sound, yet they drove significantly stronger cortical responses when self-generated despite substantial overlap in the neural populations responsive in the passive condition (60% ± 15%). This precludes the possibility that our findings are driven by other known modulatory phenomena such as stimulus-specific adaptation^27,28^.

The acoustically selective suppression of neural responses to self-generated sounds was also recapitulated at the level of individual neurons. Many individual neurons displayed strong suppression of their responses to expected sounds, but not to any other sound (Fig. 2A), suggesting a highly selective circuit mechanism for attenuating neural activity. To quantify the magnitude of this prediction-based modulation at the single-neuron level, we computed a modulation index that compared a neuron’s response to each sound heard in the active and passive conditions^11^. The majority of neurons had weaker responses to the expected sound when it was self-generated compared to when it was heard passively (negative modulation values) (Fig. 2B). In contrast, neurons displayed less suppression to all unexpected sounds (p<0.01 for all), responding equally strongly on average to probe sounds when they were self-generated and heard passively, with some neurons enhanced, some suppressed, and many cells responding equally across the two conditions. The notable exception was the frequency probe, which generated enhanced neural responses relative to the passive condition, consistent with large population-level neural responses (see Fig. 1B). The different patterns of population-level activity evoked by passive and self-generated sounds were sufficient to decode the sound identify and behavioral context in which it was heard on individual trials from small groups of auditory cortex neurons (Fig. 2C). Taken together, these data are consistent with the auditory cortex simultaneously predicting the expected frequency, position, intensity, and spatial location of a self-generated sound.

**Figure 2:**
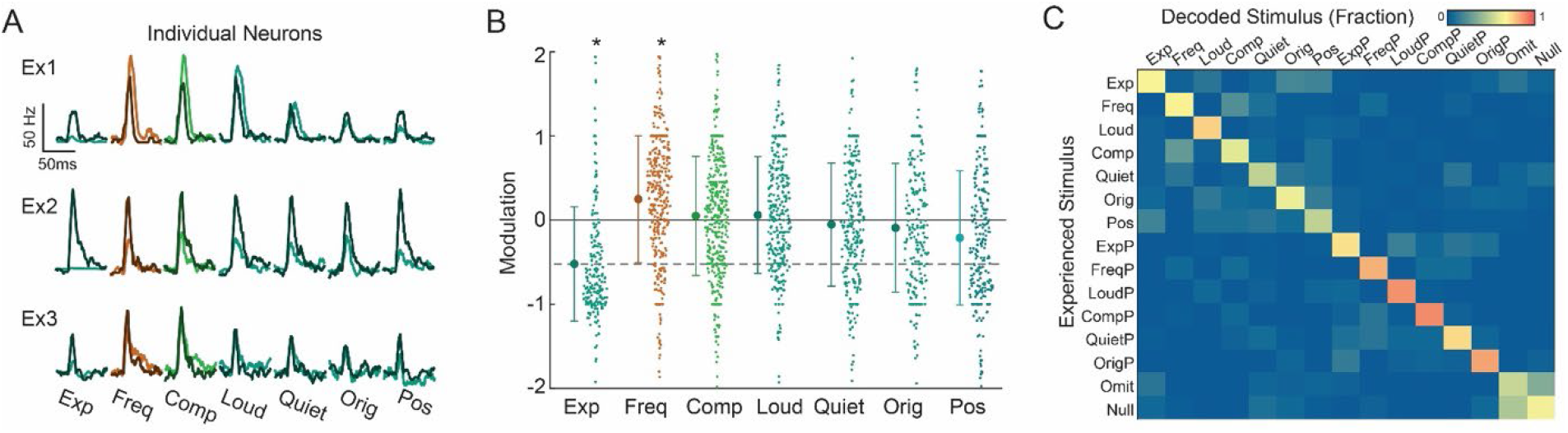
Precise suppression of expected sound responses in individual neurons. (A) Average responses across trials of three individual neurons to each tone type, showing suppression that is specific for the expected sound at the individual neuron level. (B) Modulation (See Methods) of individual neurons comparing responses to sounds heard in the active and passive condition to each tone type. Negative values indicate weaker responses in the active condition, i.e. suppression. A one-way ANOVA detected differences amongst the groups (F-statistic p = 2*10^−32^), with Exp and Freq being significantly different from all other groups (Exp, p < 1*10^−5^; Freq, p < 0.01). Neuron values and inclusion are the same as (Fig 1B). (C) Confusion matrix showing how delivered stimuli were classified from auditory cortex neural responses on individual trials. For each animal (N = 4), we measured the response of each neuron to a given sound on 20 individual trials for each sound type, omission trials, and a set of randomly selected time points during behavior that were < 0.3s away from other sounds (‘Null’). Values represent the fraction of each ground-truth trial type that were classified as a given stimulus based on neural data, averaged across 4 animals.

### Prediction error neurons respond to unexpected consequences of movement

The single-neuron analyses outlined above reveal many neurons that respond more strongly to an unexpected self-generated sound than to the same sound heard passively ^11,21,22^. While some of these neurons are likely responsive in both behavioral conditions but with relatively larger responses in the active condition, the number of strongly enhanced neurons (i.e. neurons with MI close to 1 in Fig 2B) for each unexpected sound raises the possibility that these sounds recruit a new group of cells that do not respond passively. We therefore quantified neurons that were activated by each sound in the passive condition, the active condition, or both. A relatively consistent number of neurons were responsive to each sound in the passive condition (‘Passive only’ and ‘Shared’, Fig. 3A). When mice heard the expected self-generated sound, only a small subset of passive-responsive neurons responded (‘Shared’). In contrast, when mice heard any unexpected sound, a substantially larger number of neurons responded including many neurons that were unresponsive to these same sounds heard passively (‘Active only’, p < 0.01 for all).

**Figure 3:**
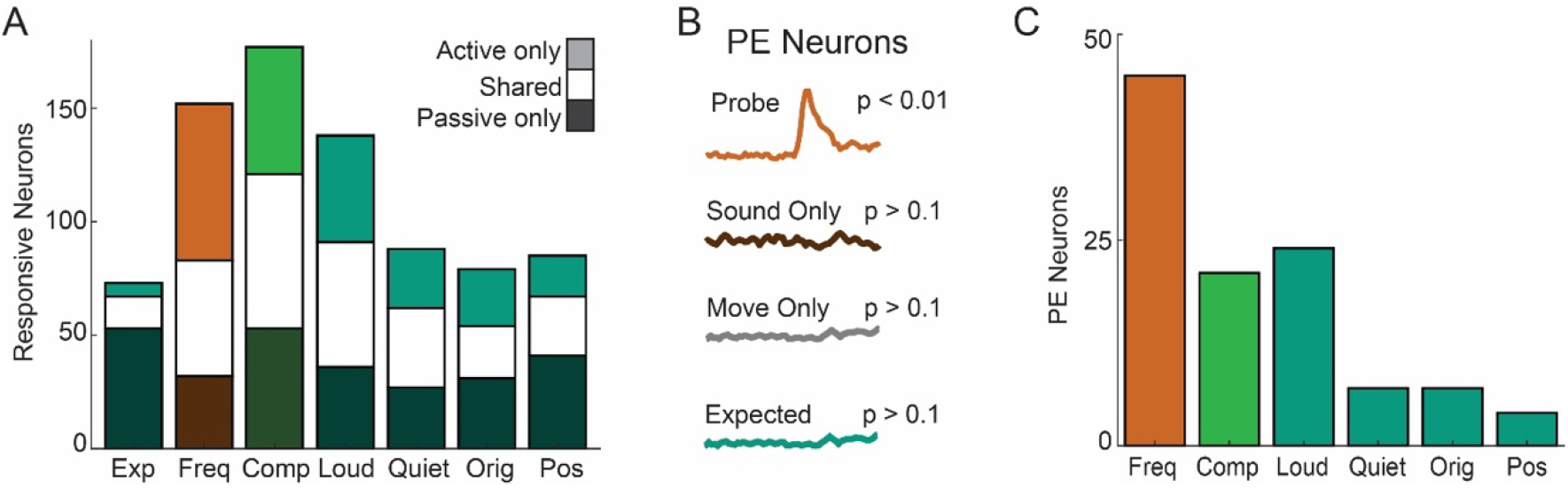
Abundant prediction error neurons in mouse auditory cortex. (A) Number of neurons responsive (p < 0.01) to a given sound in the active context (light), passive context (dark), or both (white). (B) Schematic depicting the identification of putative prediction error neurons, defined as neurons which respond to a given stimulus type in the active context, but not in the passive context, not at the time of expected sound on omission trials, and not to the expected self-generated sound. Stimulus window of 0 - 60ms post sound onset compared to the 60ms prior to sound onset. (C) Number of neurons that fulfil our putative prediction error criteria for each unexpected trial type.

Since these ‘active only’ neurons were abundantly recruited following unexpected, but not expected, self-generated sounds, we hypothesized that they may explicitly encode prediction errors. Enhanced neural responses following unexpected stimuli have been observed at the population and single-neuron level in prior experiments^11,12,16,17,22^, but it has not been conclusively established whether such responses depend upon a learned motor-sensory prediction. To determine whether prediction error neurons exist in the auditory cortex, we identify a subset of ‘active only’ neurons as putative prediction error neurons and measure their abundance following each sound in trained and untrained animals.

First, we established a stringent definition for putative prediction error (PE) neurons in the auditory cortex. We required that PE neurons respond to an unexpected self-generated sound (p<0.01) but not to the same sound heard passively, not to the expected self-generated sound, and not in the same window during silent movements (Fig 3B). This ensures that our putative prediction error neurons only respond to the presence of a sound that is self-generated and unexpected, and cannot arise due directly to movement, or to the enhancement of passive sound responses. Using these criteria, we identified 85 PE neurons, corresponding to 8.4% of all recorded neurons and 29.8% of sound-responsive neurons (Fig 3C).

Our strict criteria for PE neurons preclude the possibility that their responses to unexpected self-generated sounds arise through a simple combination of suprathreshold sound and movement responses. However, prediction error-like signals could potentially arise through subthreshold mechanisms that are unrelated to expectation but instead reflect a simple convergence of subthreshold motor and auditory inputs^25^. We reasoned that if the PE neurons we describe here truly reflect experience-dependent violations from expectation, then these neurons should not be present in mice that did not expect the lever to produce a sound. To test this, we trained a separate cohort of mice to make identical lever movements, but in silence. We recorded neural responses to a variety of self-generated and passive sounds in these silent-trained mice and found that only a very small fraction of cells fulfilled our criteria as PE neurons (Fig 4A). The comparative abundance of neurons responsive only to unexpected self-generated sounds in sound-trained mice compared to silent-trained mice demonstrate that the putative prediction error neurons identified by our criteria reflect the violation of a learned expectation.

**Figure 4:**
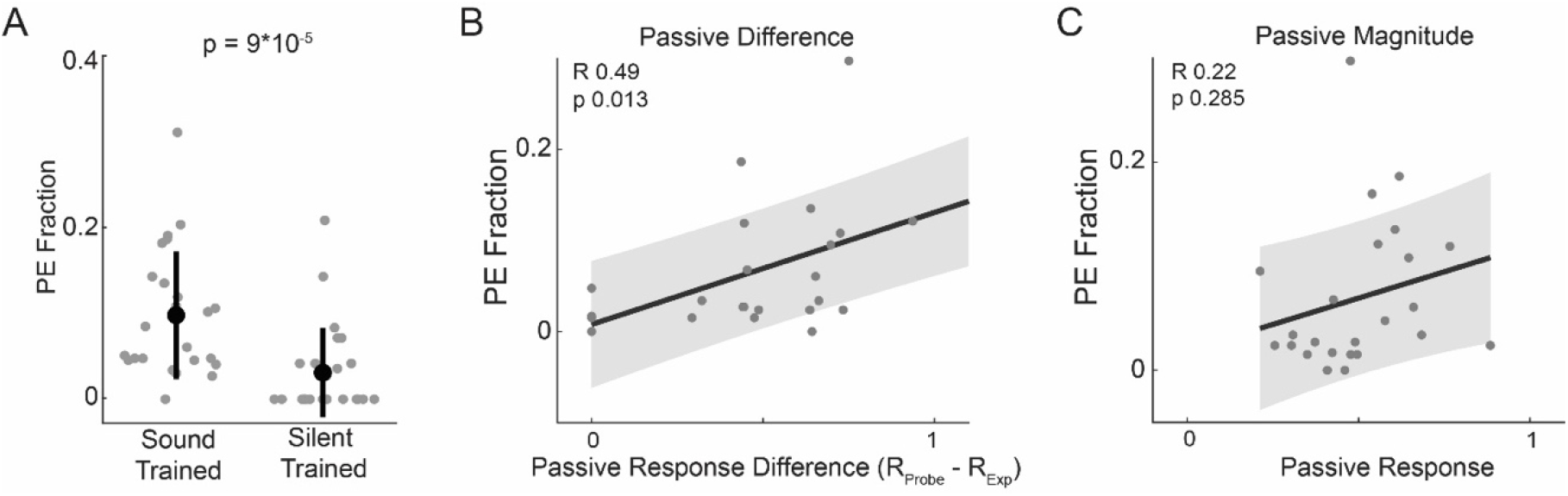
Prediction error neurons reflect the violation of a learned expectation. (A) Quantification of the number of putative prediction error neurons in trained animals, and animals trained on an identical but silent lever task. Each dot represents the fraction of total number of neurons in a recording session that met the criteria for prediction error neurons for a given stimulus. (B) Comparison between the number of prediction error neurons for a stimulus (as in A) and how ‘different’ a stimulus was from the expected sound. Differences were quantified between neural responses to each probe sound and the expected sound in the passive condition (See Methods). Each dot represents one unexpected stimulus in one animal (N = 4), and difference values were mean-normalized within animal to enable a comparison across animals. Linear regression is shown with shaded standard error. P values and correlation coefficients are listed. (C) Identical analysis as (B) but using the absolute magnitude of an animal’s population response to each stimulus heard in the passive condition.

A hallmark characteristic of prediction error neurons throughout the brain is the scaling of error responses with the magnitude of the perceived error^29,30^. Given that different probe stimuli evoked different numbers of PE neurons (See Fig 3C), we asked whether the number of PE neurons recruited by a stimulus was related to how different the stimulus was from “expected.” We measured stimulus similarity using a population-level neurometric approach, computing the absolute difference between a neuron’s response to the expected sound and a probe sound, summed across all neurons in an animal. During passive playback, some probe sounds evoked population-level responses more similar to the expected sound while other probe sounds produced highly dissimilar responses (Fig. 4B). We used this measure of response similarity relative to the expected sound as a proxy for how strongly a stimulus violated expectation. We observed that the number of PE neurons responsive to an unexpected sound scaled with the magnitude of the estimated expectation violation (Fig 4B). To ensure that this finding was not simply because some sounds activated the auditory cortex more strongly in general, we performed a similar analysis, comparing the number of PE neurons to the magnitude of a sound’s response in the passive condition (Fig 4C). We found no correlation between the number of PE neurons evoked by a sound and passive response strength, supporting the conclusion that the number of PE neurons observed reflects the ‘unexpectedness’ of a movement’s sensory outcome.

Together, these findings identify a substantial population of sensory prediction error neurons in the auditory cortex whose responses signal the violation of a learned motor-sensory expectation.

### Prediction error neurons signal specific rather than generic violations

Our definition of a PE neuron requires that it is responsive to a self-generated probe sound and unresponsive to the same sound heard passively. Neurons that fulfil these criteria could be highly selective for a single self-generated sound but could also respond to other sound types in either the active or passive condition. To determine the specificity of auditory cortex PE neurons, we visualized each neuron by displaying its responsiveness across active and passive stimuli, and the stimuli for which it signals a prediction error (Fig 5A). Auditory cortex PE neurons fell into two general categories: Neurons that responded only to one or two unexpected self-generated sounds and no passive stimuli (Fig 5B, Neuron 1), or neurons that responded to a different set of stimuli in the active and passive condition (Fig 5B, Neuron 2).

**Figure 5:**
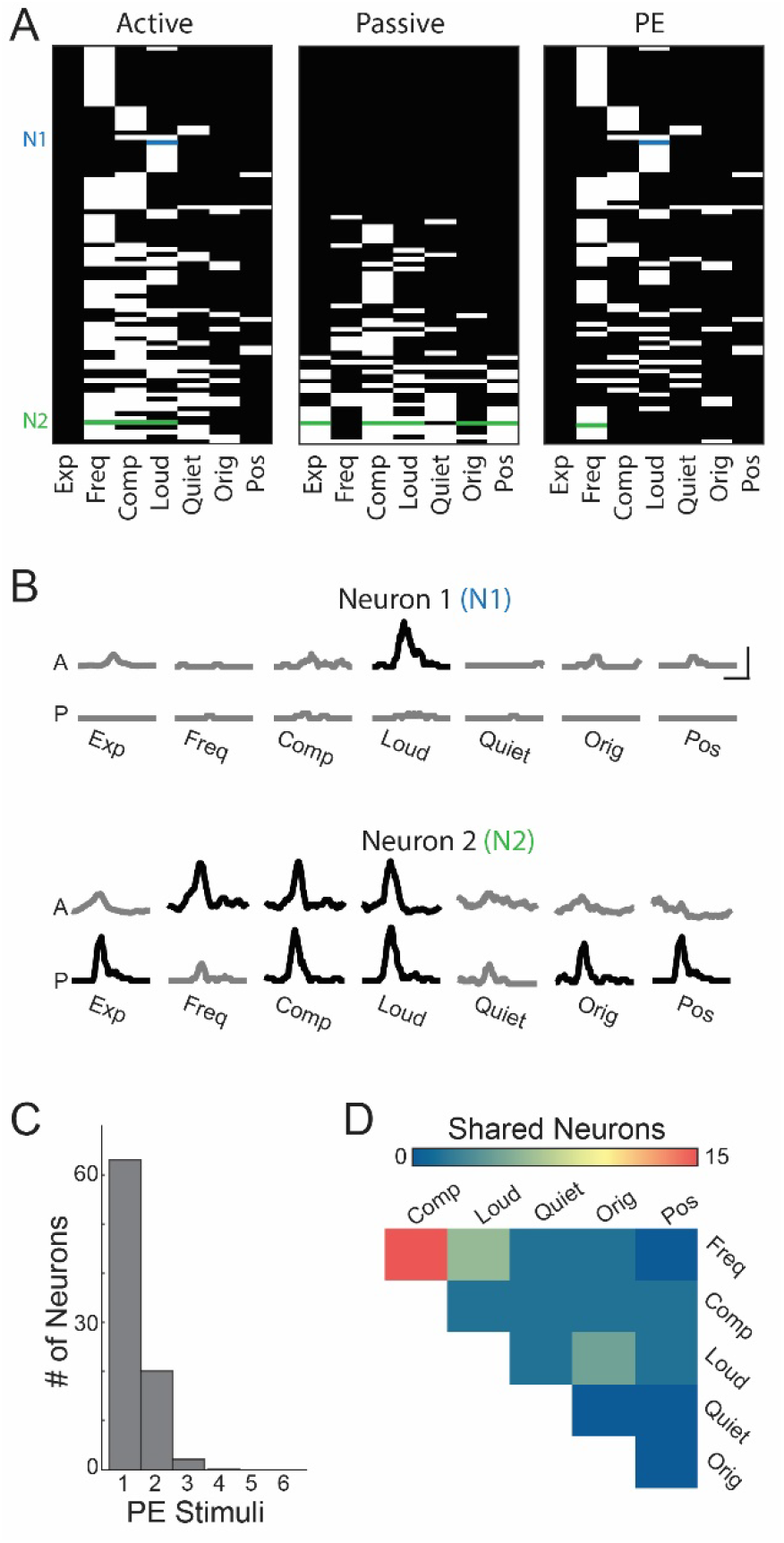
Prediction error neurons are stimulus-specific. (A) Visual representation of each prediction error neuron’s responsiveness (white) to task tones heard in the active condition (left), responsiveness in passive condition (middle), and whether a neuron obeyed our prediction error criteria for a given stimulus (right, see Fig 3B). To match our prediction error criteria, a probability value of 0.1 was used as a cutoff for the expected sound (first column in each map), while all others reflect a cutoff of 0.01. Rows with color represent example neurons in (B). (B) Responses of two example neurons to sounds heard actively (top) and passively (bottom). Black PSTHs show significant responses using the p values described in (A). (C) Quantification of the number of different stimuli for which a neuron signals prediction error. (D) Color-coded matrix showing the number of prediction error neurons that are shared across pairs of stimuli.

Nearly half of auditory cortex PE neurons (45%) were unresponsive to any of the passive sounds that we presented, with 70% responding to one or fewer, suggesting that many of these neurons would not classically be considered sound-responsive neurons. Most PE neurons signaled a prediction error for only one unexpected outcome (74%) and 97% of PE neurons signaled two or fewer outcomes, consistent with PE neurons signaling specific rather than generic errors (Fig 5C). For the subset of PE neurons that responded to multiple unexpected self-generated sounds, we evaluated the specific sets of violation stimuli by which they were activated (Fig 5D). The vast majority of these non-specific PE neurons were responsive to the frequency probe and composite probe stimuli. This pairing makes sense since both the composite and frequency probe stimuli contained the same unexpected frequency.

Stimulus-specific PE neurons could arise through computations of prediction errors at a higher cortical level that are transmitted back to auditory cortex^23^ or could arise *de novo* in the auditory cortex through a local mechanism. These local versus top-down computations should result in error signals with different latencies; top-down errors would arise later than would local computations. We therefore quantified the onset latency of prediction error neurons measured as the time to first spike following stimulus onset (Fig 6A). Error responses to the frequency probe were as rapid as neural responses to passively heard sounds and responses to self-generated sounds by non-PE neurons (Fig 6B). Given the specificity and early onset of prediction error signals following unexpected sounds, it is unlikely that these neurons are driven by feedback of a general error signal calculated elsewhere in the brain. Instead, PE neurons in the auditory cortex may reflect errors that are computed locally through the integration of bottom-up sensory input and top-down expectations.

**Figure 6:**
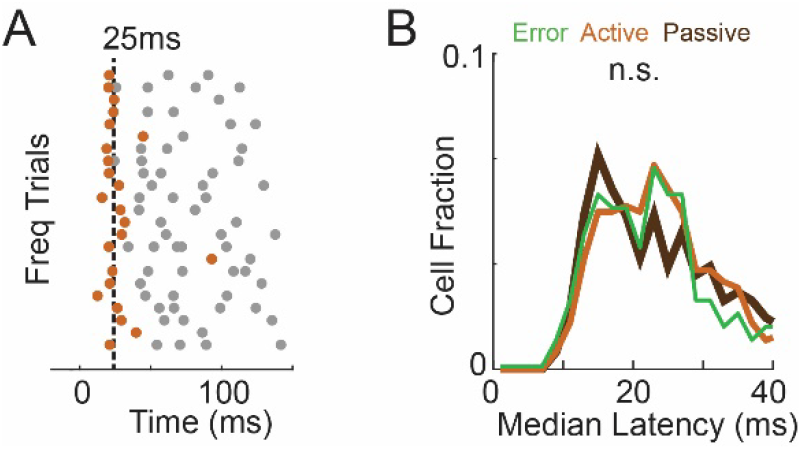
Prediction error responses in auditory cortex are short-latency. (A) Raster of example neuron showing action potential timing following frequency probe sounds, with the first spike on a given trial (orange) used to calculate an average onset latency. (B) Histogram of average onset latency following frequency probe trials for prediction error neurons (Green), all neurons responsive to the frequency probe (Orange), and latency of neurons responsive to the passive frequency probe following passive presentation. No difference between prediction error neuron latencies and general latencies in the active condition (p = 0.97) or to passive sound responses (p = 0.97, p = 0.87, KS Test).

## Discussion

Overall, our data reveal that movement-based predictions result in sound-suppression and violation-signaling that are specific across multiple feature dimensions and likely to arise de novo in the auditory cortex.

Auditory cortex activity displayed prediction-based suppression that was specific for the frequency, intensity, timing, and spatial origin of an expected sound when measured at the population activity level and when measuring individual neuron modulation. A simple circuit model in which somatic inhibition decreases the spiking of neurons tuned to the expected stimulus would lead to comparable inhibition in a given neuron for both expected and unexpected sounds^6,13,31–33^. Instead, we observed many individual neurons that experienced strong suppression of responses to the expected sound during movement while experiencing weaker suppression or even enhancement in response to other self-generated sounds (Fig 2A). These data suggest a more subtle and targeted form of inhibition that can filter neural responses to an expected sensory outcome across multiple features simultaneously within a single neuron.

We also observe abundant sensory prediction error neurons in the auditory cortex only when a movement has an unexpected acoustic outcome. Our strict criteria exclude neurons that respond to a given sound heard passively, or that respond in the absence of sound on omission trials, eliminating the possibility that our PE neurons arise from a simple combination of sensory or motor tuning. Instead, we demonstrate that neurons with a prediction error phenotype are abundant only in animals that have a learned, motor-sensory expectation, and that the number of prediction error neurons recruited by an unexpected stimulus reflects how different the stimulus was from expectation. Individual prediction error neurons typically respond with short latency and to just one or two probe stimuli, indicating that these neurons do not reflect the feedback of a generic error signal calculated elsewhere in the brain. Our findings establish the presence of stimulus-specific sensory-prediction error neurons in the auditory cortex and suggest a cortical origin for *de novo* error calculation.

## Methods

### Animals

All experimental protocols were approved by New York University’s Animal Use and Welfare Committee. Male and female wild-type (C57BL/6) mice were purchased from Jackson Laboratories and were subsequently housed and bred in an onsite vivarium. We used 2–4 month-old mice for our experiments that were kept on a reverse day-night cycle (12h day, 12 h night).

### Surgeries

For all surgical procedures, mice were anaesthetized under isolfurane (1-2% in O_2_) and placed in a stereotaxic holder (Kopf), skin was removed over the top of the head, and a Y-shaped titanium headpost (H.E. Parmer) was attached to the skull using a transparent adhesive (Metabond). Mice were treated with an analgesic (Meloxicam SR) and allowed to recover for 5 days prior to training. Following training and 24 to 48 hours prior to electrophysiology, a small craniotomy was made to expose the auditory cortex (∼2mm diameter, -2.5mm posterior, 4.2 mm left from bregma). Another small craniotomy was made above the right sensory cortex and a silver-chloride reference electrode was positioned atop the surface of the brain for use as a ground electrode and covered (Metabond). Exposed craniotomies were covered with a silicone elastomer (Kwik-Sil) and the mouse was allowed to recover in its home cage, and an additional training session was performed prior to electrophysiology.

### Behavioral Training and Data Collection

We adapted a custom head-restrained lever-based behavioral training paradigm where mice push a lever and hear closed-loop sounds^11^. A custom-designed lever (7cm long, 3D printed using Formlabs Form2) was mounted to the post of a rotary encoder (US Digital) 5cm from the lever handle. A magnet (CMS magnetics) was mounted to the bottom of the lever, which was positioned 4cm above a larger static magnet which established the lever resting position and provided light and adjustable movement resistance. The lever handle (top) was positioned adjacent to a tube (Custom, 3D printed using Formlabs Form2) to hold mice directly below two plate clamps (Altechna) to secure the mouse headpost. Lever and mouse apparatus was constructed with Thor-labs components. A water tube, controlled by a solenoid valve (The Lee Company), was positioned in front of the mouse. Digital signals for lever movement were collected by a data acquisition card (National Instruments) connected to a computer and logged by custom Matlab software (Mathworks, PsychToolBox) and sampled at 2Khz. Digital processing of lever movements received sufficient processing in real time to track important movement thresholds, which were used to trigger sound events based on user-defined closed-loop rules. Sound output was delivered from the computer to a sound card (RME Fireface UCX), the output of which was routed to an ultrasonic speaker (Tucker Davis Technologies) located lateral to the mouse, ∼10cm from the mouse’s right or left ear. We recorded sounds during test experiments using an ultrasonic microphone (Avisoft, Model # CM16/CMPA-P48) positioned 5 cm from the lever to confirm that the lever produced negligible noise (<1 dB SPL) and that experimenter-controlled sounds were delivered at a consistent volume of 50, 65, and 80 db depending on stimulus type. All training was performed in a sound-attenuating booth (Gretch-Ken) to minimize background sound and monitored in real-time via IR video.

During lever training, mice were water restricted and maintained greater than 80% of pre-restriction body weight and received all of their water (1-2ml) while performing the lever behavior. In practice, body weight was often above 90% since diminished body weight was not necessary to induce lever pressing once mice learned the task. During training, mice were head-fixed to the behavioral apparatus and presented with the lever and lick-port after ∼10 minutes of quiet acclimation. Mice were then allowed to make outwards lever movements at will. For a movement to be considered valid, we required the lever to remain in the home position (∼+/-3mm from rest) for >200 ms prior to initiation. Valid movements that reached a reward threshold (∼15mm from home position) elicited a small water reward (5-10uL) when the lever returned to home position. Auditory feedback in the form of a pure tone (50ms duration, 65dB, 12khz) was delivered on all trials when the lever crossed a set threshold 1/3 of the way between the home position and reward threshold for the first time in a trial. To ensure strong coupling between movement and sound, auditory feedback was provided on all trials, regardless of whether mice obeyed the home-position requirement and would subsequently receive a reward. Initially, 100% of successful trials produced a reward, but over the course of training that number was dropped to 25% to produce more lever movements per session. The reward rate was stable for at least 5 sessions prior to recording. Overall, mice received between 18 and 22 sessions of training over 10-12 days prior to electrophysiology, with either one or two sessions per day.

### Electrophysiological Recording and Aggregate Neural Responses

Following training, surgical opening of a craniotomy, and one subsequent training session, mice were positioned in the behavioral apparatus and a 128-channel electrode (128AxN, Masmanidis Lab) was lowered into the auditory cortex orthogonal to the pial surface^34^. The electrode was connected to a digitizing head stage (Intan) and electrode signals were acquired at 30khz, monitored in real time, and stored for offline analysis (OpenEphys). The probe was allowed to settle for at least 20 minutes, at which point the lever and lick-port were introduced and mice were allowed to make lever movements at will as in any other training session. After performing at least 30 standard lever movements, we unexpectedly began a probe session in which mice heard several different sounds. 93% of sounds were as expected (‘Exp’, 12khz, 65db) while 1% each were a substituted frequency (‘Freq’, 5.6khz, 1.1 octave lower, 65db), both the unexpected and an unexpected frequency (‘Comp’, 5.6khz and 12khz, 65db), a higher intensity (‘Loud’, 12khz, 80db), a lower intensity (‘Quiet’, 12khz 50db), played from a different origin (‘Orig’, 12khz, 65db, played from a speaker on the left side of the mouse’s head), played during the return phase of the lever movement (‘Pos’, 12khz, 65db, half way between reward threshold and the return to the home position on trials reaching reward threshold), or omitted. The requirements for reward delivery were not influenced by the identify or timing of auditory feedback. Following probe sessions, the lever was removed and tone frequencies ranging from 3 to 32kHz (0.5 octave spacing) as well as all tones presented during the active phase of the task were presented with random inter-tone intervals drawn from a flat distribution with range 1 to 2 seconds.

After recording, electrical signals were processed and the action-potentials of individual neurons were sorted using Kilosort2.5^35^, and manually reviewed in Phy2 based on reported contamination, waveform principal component analysis, and inter-spike interval histograms. Because the identification of prediction error neurons could be dramatically skewed by the loss of neural signals over the course of an experiment, we excluded any neuron that had a statistically significant difference (p<0.05) in baseline firing rate or the response rate to passively heard tones from the pre- and post-behavioral passive tone sessions. We analyzed neurons with non-fast-spiking waveforms, separated by plotting peak to valley ratio against action potential width. Tone-evoked average firing rate PSTHs were measured in 2 ms bins and aligned to sound onset for each neuron for each tone type (Fig 1B). PSTHs and individual neuron modulation for a given tone type include all neurons that were responsive (p<0.01) to a given tone in either the active or passive condition measured as an increase in firing rate from baseline (60ms prior to stimulus onset) during the sound response window (0-60ms post stimulus onset) across trials using a paired rank sum test. To measure the movement-based modulation of each neuron’s responses to the lever-associated or probe tones, we compared the neural sound response in our analysis window to the same sound in the active and passive condition using a radial modulation index^11^. Radial modulation was calculated as the theta value resulting from a cartesian to polar transformation of the response strength in the active condition compared to the response strength in the passive condition. Theta values were converted to a scale of +/-2 and rotated such that a value of 0 corresponded to equal responses across the two conditions. The fraction of neuron overlap reported in the text measures the fraction of neurons responsive to the passively heard expected sound that also respond to each probe sound.

In a subset of animals, we performed electrophysiological recording of mice that had been trained on an identical version of the lever task but without sound feedback. On experiment day mice first performed silent lever pushes for 20-50 trials, then we delivered a range of sound frequencies (4-24khz, half octave intervals, 50ms duration, 65dB,) at the sound threshold during lever pushes, followed by presentation of the same sounds passively with the lever removed, as above.

### Prediction error Neuron Analysis

We defined prediction error neurons as having a significant response in the sound response window (p<0.01, 0-60ms post stimulus onset compared to 60ms prior to stimulus onset) for a given stimulus type, but not to the same stimulus heard passively (p>0.1), to the expected sound heard actively (p<0.01) or at the same position during movement on omission trials (p>0.1). Prediction error neurons were identified independently for each stimulus type. Prediction error neurons were identified in silent trained animals using the same functional definition comparing activity in the movement condition, passive sound condition, and active condition.

The fraction of prediction error neurons (Fig 4A-C) was defined as the number of prediction error neurons for a stimulus type divided by the total number of sound-responsive neurons in an experiment (Active or passive) and are presented with data points representing one stimulus in one animal. For analyses involving individual animals (Fig2C, Fig4A-C), data was analyzed only for animals that had more than 40 sound responsive neurons in the population (N = 4). For regression comparisons, the neurometric difference between a probe stimulus and the expected stimulus was calculated by comparing average response responses across the two tones in the passive condition. The difference between responses to the two tones for each neuron was summated across all neurons in an animal and used to represent the dissimilarity of neural response patterns between the probe sound an expected sound. These values were mean normalized within each animal to allow for comparison across animals. A similar process was used for passive response magnitude, but with average firing rates summated across all neurons in an animal instead of making a comparison to the expected sound. Onset latencies were defined for each neuron as the average of first post-stimulus spike times on each trial. Trials that did not produce an action potential in the sound response window were removed from the average. Histograms of onset latencies were created using 2ms bins.

### Decoding Analysis

Decoding data were organized in a trials-by-neuron matrix within each animal, with each cell representing the response of an individual neuron on an individual trial. A consistent number of trials (20, randomly selected) was used for each stimulus type. Each trial, in sequence, was removed from the data set, and the remaining trials along with the ground truth identity of the experienced stimulus was used to train a multiclass error-correcting output codes model using support vector machine binary learners^36,37^. The trained model was then used to classify the withheld trial, which was then compared to the ground truth identify of the stimulus. This process was repeated for all trials in an animal, with the results visualized as a confusion matrix comparing the classification result to the ground truth identity of each trial. Each pixel represents the number of trials classified as a given stimulus type divided by the number of ground truth trials for a given stimulus type. The resultant confusion matrices were then averaged across animals.

### Statistical Analysis

Throughout, animal values are denoted by a capital N while cell values are denoted by a lowercase n. Unless otherwise reported, all averages and error bars denote mean ± standard deviation. P values are reported in text or on the relevant figure panels for all statistical comparisons. Statistical comparison of aggregate neural activity use a one-way ANOVA followed by two-sided, non-paired, non-parametric rank sum test and Bonferroni correction for multiple comparisons. The comparison of the number of ‘active only’ neurons for probe stimuli vs. the expected stimulus was performed by bootstrap resampling, with which we compared the observed counts for the two stimuli to 10,000 randomly generated distribution of counts created assuming equal probability. Statistical comparison of onset latency across groups was performed using a Kolmogorov-Smirnov test. The relationship between the number of prediction error neurons and neural response properties was measured using linear regression and correlation coefficient analysis with p and R values reported.

## Acknowledgements

We thank Alessandro La Chioma, Ralph Peterson, and Grant Zempolich for their thoughtful comments on the manuscript. We thank members of the Schneider lab for fruitful discussions. We thank Jessica A Guevara for expert animal care and technical support. This research was supported by the National Institutes of Health (1R01-DC018802 to DMS); a Career Award at the Scientific Interface from the Burroughs Wellcome Fund (D.M.S); fellowships from the Searle Scholars Program, the Alfred P. Sloan Foundation, and the McKnight Foundation (D.M.S.); and an investigator award from the New York Stem Cell Foundation (D.M.S). D.M.S. is a New York Stem Cell Foundation -Robertson Neuroscience Investigator.

## Declaration of Interests

The authors declare no competing interests.

## Notes

### Competing Interest Statement

The authors have declared no competing interest.

### Summary of Updates

Minor analysis and figure changes

